# Just add water: A simple floral bud injection method for stable *Agrobacterium*-mediated transformation in two ecotypes of *Mimulus guttatus*

**DOI:** 10.1101/2024.06.24.600443

**Authors:** Lauren E. Stanley, David B. Lowry

## Abstract

Stable transformation is the biggest challenge for studying gene function in plants. In most plant species, stable transformation requires the development of arduous tissue culture and regeneration methods that often fail and can cause undesirable genetic changes. Floral transformation methods bypass these challenges by infiltrating *Agrobacterium* directly into the ovules of developing flowers. However, this method has had limited success outside of a handful of plant species and genotypes within those species. Here, we demonstrate that the floral infiltration method is effective for some, but not all genotypes of the yellow monkeyflower, *Mimulus guttatus*, a model system for studies of ecology and evolution. To conduct transformation in genotypes where infiltration failed, we developed a novel floral injection method. Beyond expanding the number of genotypes that can be transformed, the floral injection method has other advantages, such as allowing plants to be infiltrated multiple times and a reduction in floral abscission and male sterility. Through a combination of floral infiltration and injection methods, we were able to transform both coastal perennial and inland annual genotypes of *M. guttatus*, which sets the stage for understanding the molecular genetic underpinnings of local adaptation to the divergent habitats occupied by these distinctive ecotypes. Overall, the development of these efficient transformation methods will open up a wide range of new research avenues in the *M. guttatus* species complex.

## INTRODUCTION

Stable transformation is the greatest single bottleneck in studying gene function in plants (Altpeter et al., 2016). There are few plant systems for which efficient transformation protocols have been developed, which greatly impedes the potential for hypothesis testing in agricultural and natural systems. While researchers can circumvent these difficulties by testing a candidate gene’s function in closely related heterologous systems, the results from such experiments should be interpreted with caution, as (epi)genomic and evolutionary context matter (Kramer, 2015). Future advances in both fundamental and applied research will rely on our ability to improve the efficiency of stable transformation in plants.

There are two major classes of stable transformation techniques currently available for plants: particle bombardment/biolistics and pathogen-mediated methods. Particle bombardment forcefully delivers DNA to plant cells in tissue culture with high-velocity gold or tungsten particles coated with the desired genetic construct (e.g., Liu et al., 2019). In contrast, pathogen-mediated methods use a disarmed pathogen, typically *Agrobacterium*, to deliver and integrate a transfer-DNA fragment either to de-/undifferentiated cells (tissue culture) or developing ovules (floral dip) (for an overview, see Hwang et al., 2017).

The vast majority of plant species require tissue culture for stable transformation, whether using particle bombardment or pathogen-mediated methods (Altpeter et al., 2016). Going through a tissue culture stage adds major challenges, as conditions have to be highly customized to produce totipotent cells in sterile culture that are transformable. Further, developing methods to regenerate whole plants from tissue culture can be unreliable and highly genotype-specific, requiring extensive expertise. The whole process is also highly time and labor intensive, potentially expensive, and can cause undesirable genetic changes during regeneration (Anjanappa & Gruissem, 2021; J. Liu et al., 2019; Long et al., 2022).

An alternative method to tissue culture is floral dip, the method that has made *Arabidopsis* a genetic powerhouse (Clough & Bent, 1998; Zhang et al., 2006). This method involves dipping a plant at the flowering stage into *Agrobacterium* resuspended in sucrose (a carbon source for the bacteria), Silwet-L77 (a surfactant that allows bacteria to better penetrate plant tissues), and, optionally, acetosyringone (a phenolic compound produced during plant wounding that increases bacterial virulence). The dipped plant is allowed to set seed and seedlings are then screened for the presence of the transgene/s. Unfortunately, this method does not work in many plant species, or is so inefficient compared to other methods that it has not been widely adopted (e.g, rice and wheat (Ratanasut et al., 2017; Zale et al., 2009)). As a result, floral dip is not used widely outside of the Brassicaceae (with some exceptions, like *Setaria* (Martins et al., 2015; Saha & Blumwald, 2016; Van Eck, 2018), lisianthus (Fang et al., 2018), *Camelina* (X. Liu et al., 2012), and *Mimulus* (see below)).

In this paper, we report the successful implementation of the floral dip method in the yellow monkeyflower (*Mimulus guttatus*; syn. *Erythranthe guttata*). We also report the development of a new floral bud injection method for transformation that overcomes some of the challenges associated with floral dip. This work builds on the successful development of a floral dip/spray transformation method combined with vacuum infiltration, which is used in other species of monkeyflower (*M. lewisii*, *M. verbenaceus*, *M. parishii, M. luteus*; described in Yuan et al., 2013, dx.doi.org/10.17504/protocols.io.3tkgnkw; based on Bechtold & Pelletier, 1998; Chung et al., 2000). Recent studies in these systems have improved our understanding of the genetics, development, and evolution of traits that are difficult or impossible to study in traditional model plant species (Yuan, 2019). These studies would not be possible without efficient transformation protocols.

We were motivated to develop more efficient methods for transformation in *M. guttatus* because of its established role as an excellent system for the study of a vast array of ecological and evolutionary processes (reviewed in Twyford & Friedman, 2015; Wu et al., 2008). The development of efficient transformation protocols for this system should open up the possibility of studying many of these ecological and evolutionary processes at the mechanistic level. Of particular interest is understanding the genetic basis of local adaptation and reproductive isolation, which is why we focused our effort here on the evolutionary divergent coastal perennial and inland annual ecotypes of *M. guttatus* (Hall et al., 2006; Hall & Willis, 2006; Lowry et al., 2008, 2019; Lowry & Willis, 2010).

Coastal perennial and inland annual populations have drastically different life-history strategies despite frequently occurring in close (<10km) geographic proximity (Hall et al., 2006; Hall & Willis, 2006; Lowry et al., 2008, 2019). The coastal perennials experience high herbivory and exposure to oceanic salt spray, while the inland annuals face a short growing season ending in terminal drought (Lowry et al., 2009; Lowry & Willis, 2010; Popovic & Lowry, 2020). These ecotypes have been shown to be locally adapted to their particular environments, with coastal perennials investing in robust vegetative growth and defense, and inland annuals devoting resources to rapid growth and reproduction (Hall & Willis, 2006; Lowry et al., 2008; Lowry & Willis, 2010). The coastal perennial/inland annual system is therefore an excellent model for studying local adaptation.

Recent work combining Quantitative Trait Locus (QTL) mapping, population genomics, and gene expression analyses has revealed a set of candidate genes underlying the divergence of coastal perennial and inland annual ecotypes (Gould et al., 2017, 2018; Lowry & Willis, 2010). While these genes have been identified, transgenic confirmation of causality and gene function has been difficult due to the lack of efficient stable transformation protocols for the system. Prior to the work we report here, transformation in the *M. guttatus* species complex was conducted using more arduous tissue culture and plant regeneration methods (e.g., Ding et al., 2020). All previous attempts to adapt a floral dip method for this species complex had been unsuccessful and remained unpublished.

In this paper, we describe a modified floral dip method and introduce a novel floral bud injection/“jab infiltration” method for stable transformation of coastal perennial and inland annual ecotypes of *Mimulus guttatus*. These methods will allow us to begin investigating the genetic mechanisms of local adaptation in our study system. We hope that this method can also be widely used by *Mimulus* researchers and complement the existing transformation methods in the group to broaden the number of genotypes amenable to transgenic manipulation.

## METHODS AND RESULTS

### Selection of plant materials

The initial goal of our research was to transform the two inbred lines used extensively in the past to study the genetic basis of local adaptation to coastal and inland habitats: the coastal perennial SWB-S1 and the inland annual LMC-L1 inbred lines. Both lines were originally derived from populations in Mendocino County, California (Friedman et al., 2015; Gould et al., 2018; Kollar et al., 2023; Lowry et al., 2019; Lowry & Willis, 2010). SWB-S1 plants (and other coastal perennials) have many features that make them amenable to floral dip transformation, such as the ability to produce many flowers simultaneously, no reduction in fertility due to inbreeding, and making many seeds per fruit (up to 1000). By contrast LMC-L1, and other inland annuals, have many features that make them less amenable to floral dip transformation, such as fewer concurrent flowers, reduced fertility due to inbreeding depression, and few seeds per fruit (up to 50). Because of major challenges that we faced attempting to transform LMC (described below), we also attempted transformation in five additional inland annual accessions: OCC, OAE, MOR (Sonoma County, CA), and SWC (Lane County, OR). Information for all accessions can be found in Table 1.

**Table 1.**
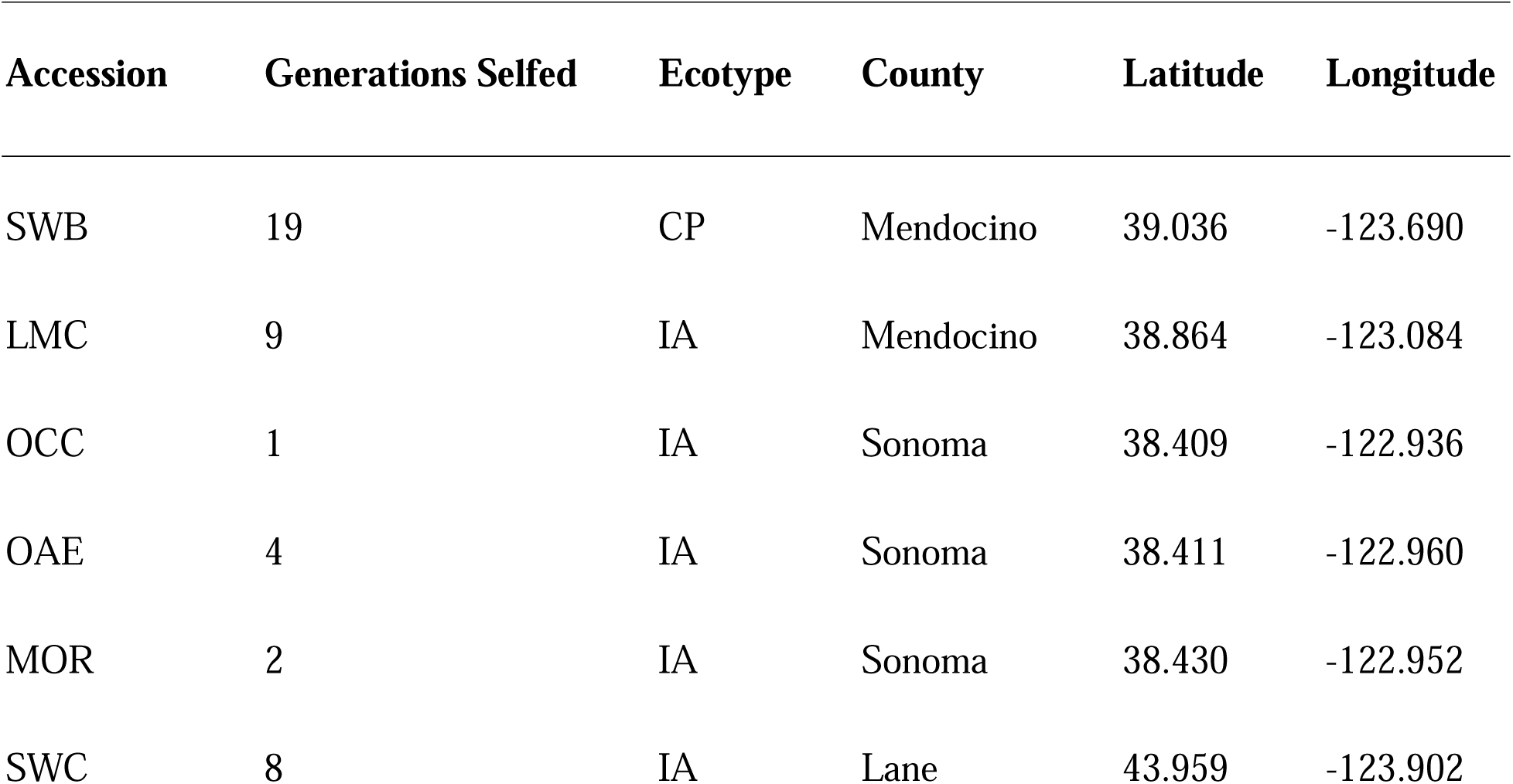
*Mimulus guttatus* accessions. All accessions used in this study, with accession, number of generations selfed, ecotype, and county and coordinates of origin (IA: inland annual, CP: coastal perennial).

| Accession | Generations Selfed | Ecotype | County | Latitude | Longitude |
| --- | --- | --- | --- | --- | --- |
| SWB | 19 | CP | Mendocino | 39.036 | -123.690 |
| LMC | 9 | IA | Mendocino | 38.864 | -123.084 |
| OCC | 1 | IA | Sonoma | 38.409 | -122.936 |
| OAE | 4 | IA | Sonoma | 38.411 | -122.960 |
| MOR | 2 | IA | Sonoma | 38.430 | -122.952 |
| SWC | 8 | IA | Lane | 43.959 | -123.902 |

### Vectors

To test transformation efficiency, the same vector was used for many infiltrations: *35S:YFP-NEGAN*. This pEarleyGate 104 construct (Earley et al., 2006) contains a 5’ Yellow Fluorescent Protein (YFP) tag and the full-length coding sequence of *NECTAR GUIDE ANTHOCYANIN* (*NEGAN*) driven by the strong CaMV 35S promoter. NEGAN (also known as MgMYB5 in *M. guttatus* and PAP2 (AT1G66390) in *Arabidopsis*), is an R2R3-MYB from *Mimulus lewisii* that activates the production of anthocyanins (Yuan et al., 2013), and thus transgenic plants can be easily identified by both resistance to the herbicide BASTA and ectopic production of purple pigment.

We have also transformed both coastal perennial and inland annual plants with other constructs for ongoing projects: knockdown RNAi and knockout CRISPR/Cas9 constructs targeting *Gibberellic Acid Insensitive (GAI)* (pB7GWIWG2(I)-*GAI*, pRGEB32-AtU6.29-BASTA-*GAI*, respectively); a knockout CRISPR/Cas9 construct targeting *GIBBERELLIN 20 OXIDASE 2 (GA20ox2)* (pYLCRISPR-*GA20ox2*); and an overexpression pEarleyGate101 *GA20ox2* construct (*35S:GA20ox2-YFP*). Vector backbones are described in Earley et al., 2006; Hunter, 2021; Karimi et al., 2002; and Ma et al., 2015.

### Floral dip in the coastal perennial SWB-S1

We based our initial approach on the efficient stable floral dip transformation method that has been successful in other *Mimulus* species (Yuan et al., 2013, dx.doi.org/10.17504/protocols.io.3tkgnkw; based on Bechtold & Pelletier, 1998; Clough & Bent, 1998). We performed this protocol in 10 mature SWB-S1 plants, whose apical meristems had been removed at flowering to encourage branching. Each plant had four to six flowering branches. Briefly, the *Agrobacterium* strain GV3101 containing *35S:YFP-NEGAN* was grown in liquid culture and then resuspended in a medium containing water, 5% sucrose, 0.1% Silwet L-77, and 0.1 M acetosyringone. The bacterial resuspension was then sprayed onto developing buds and plants were placed in a vacuum chamber, where the pressure was held at -28 inHg for two minutes. Plants were kept at high humidity overnight, then self-pollinated for two weeks, and seeds were collected and screened on soil using 1:1000 BASTA/Finale (glufosinate-ammonium). Unfortunately, this initial protocol caused significant necrosis to flowering branches, preventing many buds from developing and causing marked male sterility, where anthers produced reduced or no pollen. No transgenics resulted from this initial transformation.

To reduce adverse effects of the *Agrobacterium*, we performed another *35S:YFP-NEGAN* transformation for six mature SWB-S1 plants, where we reduced the concentration of the *Agrobacterium* by 20%, and the concentration of the Silwet L-77 to 0.08%. This altered protocol resulted in two transgenics that were resistant to the herbicide and produced anthocyanin-pigmented vegetative and floral tissues (Figure 1).

**Figure 1.**
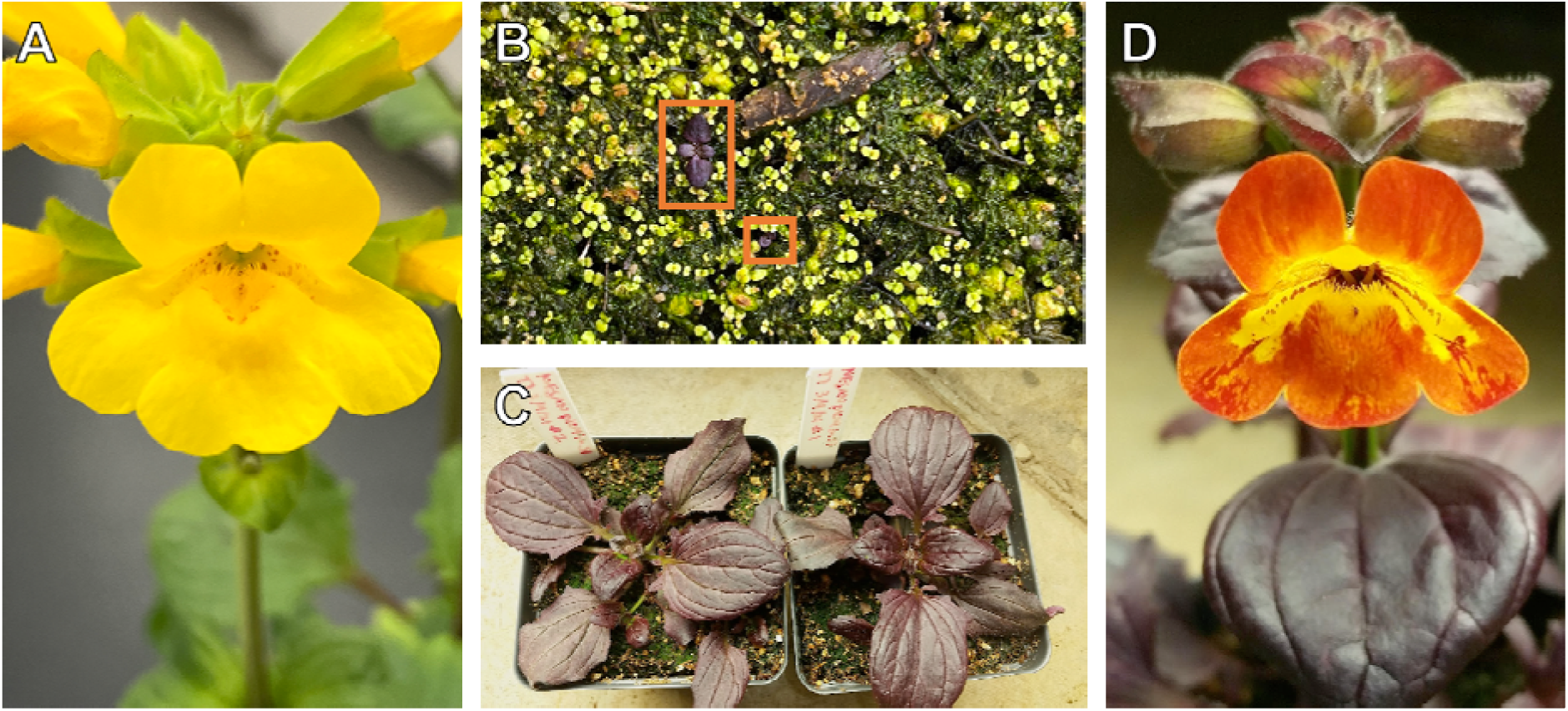
35S:YFP-NEGAN transgenics in the SWB-S1 background. **(A)** Wild-type SWB-S1 flower. **(B)** Screening for transgenics on soil. Transgenic plants are purple and indicated by orange boxes. **(C**) Young 35S:YFP-NEGAN SWB-S1 plants showing vegetative phenotype. **(D)** Floral phenotype of a 35S:YFP-NEGAN SWB-S1 plant.

We also tested the effects of life history stage on transformation efficiency. Using the modified floral dip protocol, we infiltrated ten young SWB-S1 plants (apical meristems intact). After spraying, the young SWB-S1 plants showed a necrosis phenotype similar to that observed in the original unmodified protocol, and did not produce any transgenics, indicating that life stage is crucially important for floral dip in this ecotype.

We have successfully used the modified floral dip protocol in other transformations for ongoing projects with different constructs in the SWB-S1 background. These include 23 transgenic lines from six plants infiltrated with the *GAI* RNAi construct, 35 transgenic lines from six plants infiltrated with the *GAI* CRISPR construct, and 99 lines from seven plants infiltrated with the *GA20ox2* over-expression construct.

### Floral dip in the inland annual LMC-L1

Using the modified floral dip protocol, we infiltrated ten mature LMC-L1 plants, whose main stems had been trimmed to encourage branching. LMC-L1 plants appeared unaffected by floral dip, with no visible necrosis and only a few undeveloped flowers. However, no transgenics were produced. We then repeated this experiment with six LMC-L2 plants (a different inbred line from the same population). These plants were similarly unaffected by agroinfiltration, and produced no transgenics. After five attempts, we concluded that the LMC accessions were not ideal for floral dip transformation.

### Jab infiltration

Because the floral dip method was not efficient enough to produce transgenics in the LMC background, we attempted to place the bacteria in closer proximity to ovules by injecting the bacteria into developing floral buds with a needle and syringe. This method was inspired by transient assays into strawberry receptacles (Hawkins et al., 2016; Hoffmann et al., 2006) and tomato fruits (Orzaez et al., 2006), where *Agrobacterium* solutions are injected directly into the developing organ.

This method was also informed by transient assays in *Mimulus* (Ding & Yuan, 2016). In the time since that paper was published, we have found that acetosyringone, sucrose, and even Silwet L-77 are unnecessary for the transient expression of transgenes in *Mimulus* leaf epidermal cells. In fact, these components of the resuspension medium cause tissue damage that negatively affects visualization of protein localization. Resuspension of bacteria in water alone is sufficient for construct expression in *Mimulus lewisii* leaf epidermal cells.

We first injected both LMC-L1 and SWB-S1 buds with *Agrobacterium* containing *35S:YFP-NEGAN* resuspended in water (and a water-only control) using a 30 Gauge needle to see if the delivery of the bacteria directly to ovules or the damage caused by injection would cause buds to abort or be infertile. When this was shown not to be the case, we performed our first jab infiltration by injecting the bases of developing flower buds with *Agrobacterium* containing *35S:YFP-NEGAN*. We injected the buds of five LMC-L1 plants and one mature SWB-S1 plant with the bacterium resuspended in water only, and five more LMC-L1 plants with the modified floral dip resuspension solution (5% sucrose, 0.08% Silwet L-77, and 0.1 M acetosyringone). All plants had four to six flowering branches. In both treatments, injection sites and ovaries turned purple four to five days after injection due to somatic transformation and expression of the construct. The flowers were then self-pollinated and seeds were screened for the presence of the transgene. From a single SWB-S1 plant, 27 transgenic lines were obtained. No LMC-L1 transgenic lines were obtained.

We then repeated the experiment several times with various resuspension solutions in both backgrounds. The results of all jab infiltration experiments are recorded in Table 2, and our suggested protocol may be found in Supplemental File 1. In brief, in the SWB-S1 background, transgenics were obtained with every attempted resuspension solution (water only, water + 5% sucrose + 0.08% Silwet L-77 + 0.1 M acetosyringone, and water + 0.01% Silwet L-77). Plants produced many more viable flowers and fruits when water only or water + 0.01% Silwet was used as the resuspension solution than when the modified floral dip solution was used. In addition, flowers injected with the modified floral dip solution showed marked male sterility, particularly in buds that were less developed at the time of injection. An untreated plant from the same inbred line was required as a pollen donor for these plants. This indicates that either sucrose or a high concentration of Silwet L-77 interferes with flower development. However, as expected, the number of transgenics produced per viable fruit is higher using the modified floral dip solution.

**Table 2.** Results of all floral bud jab infiltration transformations. All floral bud injection transformations carried out in this study, in chronological order. The accession and ecotype of plants used, components of the bacterial resuspension solution, the number of plants infiltrated, the number of fruits collected, and the number of transgenics obtained are provided. (CP: coastal perennial, IA: inland annual; -: information not recorded).

| Accession | Ecotype | Resuspension solution | Plants | Fruits | Transgenics |
| --- | --- | --- | --- | --- | --- |
| SWB | CP | Water | 1 | - | 27 |
| LMC | IA | Water | 5 | - | 0 |
| LMC | IA | Water, 5% sucrose, 0.08%<br>Silwet L-77, 0.1 M<br>acetosyringone | 5 | - | 0 |
| SWB | CP | Water, 5% sucrose, 0.08%<br>Silwet L-77, 0.1 M<br>acetosyringone | 3 | 41 | 102 |
| SWB | CP | Water | 3 | 118 | 31 |
| SWB | CP | Water, 0.01% Silwet L-77 | 2 | 51 | 34 |
| LMC | IA | Water | 16 | 518 | 0 |
| OCC | IA | Water, 2.5% sucrose | 9 | - | 0 |
| OCC | IA | Water, 10% sucrose | 8 | - | 0 |
| OAE | IA | Water, 2.5% sucrose | 8 | - | 2 |
| OAE | IA | Water, 10% sucrose | 8 | - | 1 |
| OAE | IA | Water | 4 | 113 | 2 |
| OAE | IA | Water, 0.01% Silwet L-77 | 4 | 112 | 1 |
| OAE | IA | Water, 2.5% sucrose, 0.01%<br>Silwet L-77 | 3 | 75 | 0 |
| MOR | IA | Water | 7 | 212 | 0 |
| MOR | IA | Water, 2.5% sucrose, 0.01%<br>Silwet L-77 | 6 | 131 | 0 |
| OAE | IA | Water | 8 | 80 | 1 |
| SWC | IA | Water | 7 | 49 | 0 |

In the LMC-L1 background, neither water-only nor modified floral dip resuspension solutions yielded successful transformations, despite the high number of buds injected. The modified floral dip resuspension solution caused most flowers to abort, while almost all buds in the water-only experiments developed (even showing somatic transformation, Figure 2). The lack of transgenics in the water-only experiments is likely due to poor seed set, which is unfortunately typical for selfed flowers in this line.

**Figure 2.**
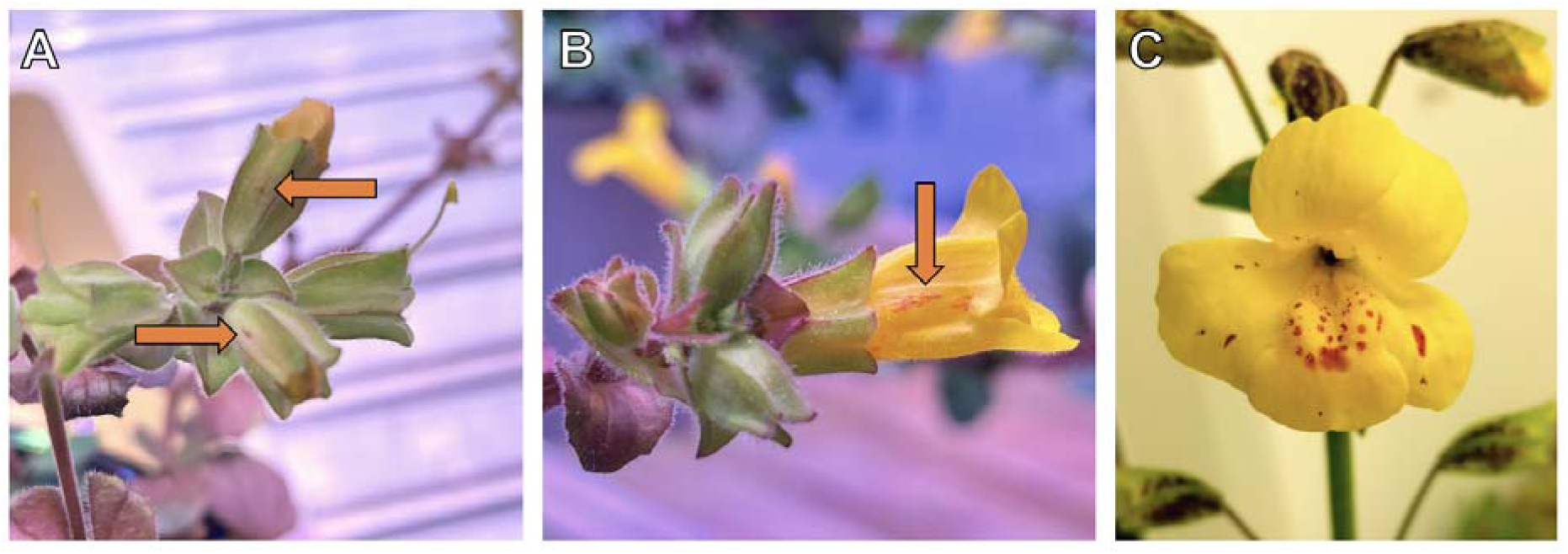
Phenotypes in T0 plants after injection with 35S:*YFP-NEGAN*. **(A)** Injection sites in SWB-S1 buds. **(B)** Somatic transformation of corolla cells near injection site in SWB-S1. **(C)** Somatic transformation in corolla of LMC-L1.

**Figure 3.**
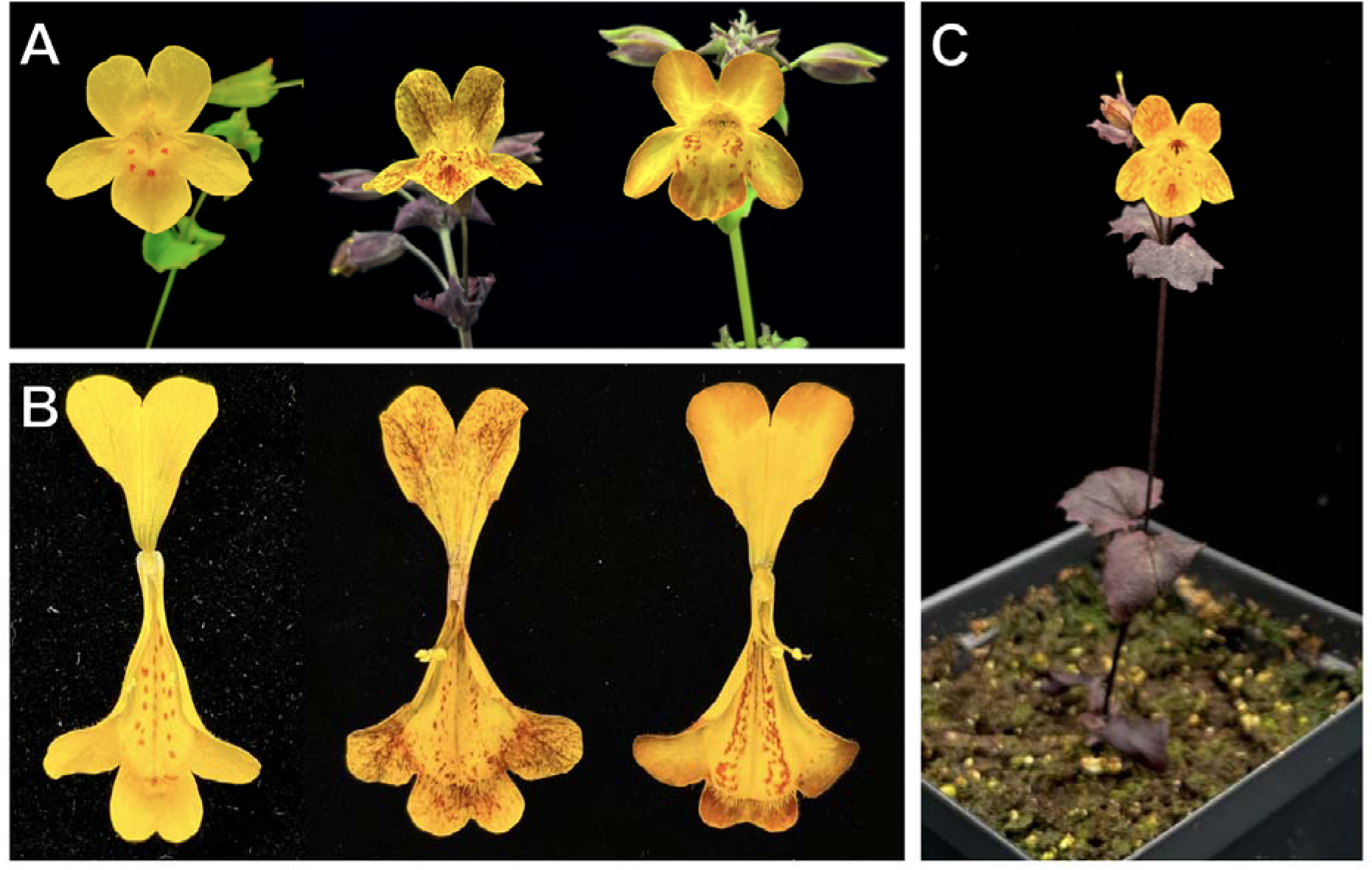
35S:*YFP-NEGAN* in the inland annual OAE background. **(A)** Front view of wild-type OAE (left) and two independent 35S:*YFP-NEGAN* lines (center and right). **(B)** Dissected corollas of wild-type OAE (left) and two independent 35S:*YFP-NEGAN* line (center and right) to show interior. **(C)** Whole plant phenotype of a third independent 35S:*YFP-NEGAN* line.

### Identification of amenable inland annual lines for transformation

Because the inland annual LMC accession was not amenable to either floral dip transformation or jab infiltration due to its low fecundity, we decided to try other inland annual accessions. These included OCC, OAE, MOR, and SWC (Table 1). For the initial jab transformation in OCC and OAE, two different resuspension solutions were tried, with eight to nine plants per line per treatment: 2.5% sucrose and 10% sucrose. No transgenics were recovered in the OCC background, but three total transgenic lines were recovered for OAE: two from the 2.5% sucrose solution experiment and one from the 10% sucrose solution experiment.

Bolstered by this result, and looking to further increase the transformation efficiency in the OAE background, we did another round of transformations in that background, with three to four plants per resuspension solution: water only, water + 0.01% Silwet L-77, and water + 2.5% sucrose + 0.01% Silwet L-77. The water-only transformation gave two transgenics from 113 fruits, the water + 0.01% Silwet L-77 transformation gave one, and the water + 2.5% sucrose + 0.01% Silwet L-77 gave none.

We also decided to try other inland annual accessions. MOR is a serpentine-adapted accession that occasionally self-pollinates in nature, which we thought might increase its chances of being amenable to transformation. Despite injecting 212 fruits with a water only resuspension solution and 131 fruits with a water + 2.5% sucrose + 0.01% Silwet L-77 solution, no transgenics were produced.

For another project in the lab we transformed a different construct, pYLCRISPR-*GA20ox2,* into two different inland annual accessions using the water-only solution: SWC and OAE. Injection of 49 fruits on seven plants resulted in no transgenics in the SWC background, but the OAE background produced one transgenic (from injecting 80 fruits on eight plants). For this particular experiment the OAE plants were in suboptimal condition at the time of injection, and still produced one transgenic plant.

## DISCUSSION

We report here successful implementation of the floral dip method in members of the *M. guttatus* species complex, a model system for understanding ecological and evolutionary processes. In addition, we developed a new floral bud jab infiltration method for *M. guttatus* that overcomes some of the disadvantages of the floral dip method. The efficiency of both of these transformation methods will greatly accelerate the testing of functional molecular genetic hypotheses in this species as well as comparative studies across this family of plants. Crucially, the jab infiltration method works in both coastal perennial and inland annual lines, which will allow us to reciprocally test gene function in genetic backgrounds that are locally adapted to contrasting environments.

### The prospects for floral bud jab infiltration

We found that transformation is feasible in both ecotypes of *M. guttatus*. Coastal perennials can be reliably transformed by both a modified floral dip protocol and a floral bud injection/jab infiltration method with several different resuspension solutions. Inland annuals are much more recalcitrant to both transformation methods, but transgenics can be obtained in certain accessions using the jab infiltration method. The modified floral dip method described here has been shown to work in an inland annual accession, SOD, from Napa County, CA (Liang et al., 2023).

Stable transformation via floral bud injection methods have been tried in other systems. Sharada et al. (2017) attempted agroinfiltration via needle injection into developing floral buds in tomato. However, this proved to be very labor-intensive and produced very few transgenics: out of 3,400 tomato floral buds injected, only 14 fruits producing 203 seeds were obtained due to floral bud abscission. There are scattered reports of similar methods working in other systems (e.g., *Paphiopedilum* orchids (Luo et al., 2021), chickpea (Shahid et al., 2016), and *Catharanthus roseus* (Bahari et al., 2020)), but these do not appear to have been widely adopted in these species, suggesting that the methodology may not be efficient for those systems. In all of these studies, researchers used standard *Arabidopsis* transformation solutions or similar, with either sucrose or glucose and acetosyringone. In all of these systems, floral damage or low seed set was reported. Based on our results, one of the key reasons that our floral bud injection method works in *Mimulus* is that the resuspension solution for the *Agrobacterium* is less harsh. Sucrose tends to be particularly damaging, and can be excluded entirely with our method. Making this modification to protocols in other systems may improve transformation efficiency in floral bud injections for other species. However, we cannot rule out that *Mimulus* is just particularly amenable to this method.

There are multiple advantages of the floral bud injection method. First, this method is much less labor-intensive than agroinfiltration through tissue culture, and avoids several issues associated with that method (e.g., contamination, somaclonal mutations). Second, it is extremely cost-effective. Floral bud injection uses very few reagents at low concentrations, and does not require expensive equipment such as the vacuum chamber and strong vacuum pump required for *Mimulus* floral dip. These two advantages make the floral bud injection method accessible across a wide range of laboratory and classroom settings.

One of the major improvements of floral bud injection over other methods is that it is less damaging to plants. This is important for two main reasons: (1) plants can be infiltrated multiple times as flower buds develop, and (2) infiltration causes less floral abscission and male sterility, particularly when using the water-only resuspension solution. These factors are especially important for genotypes or accessions that do not produce many concurrent flowers, have low pollen fertility, or have low seed set. It also circumvents the issue of plants needing to be a certain age at the time of infiltration, which is crucial for floral dip in our SWB-S1 plants and in other *Mimulus* species. The jab infiltration method could also be effective for species that have well-defended buds, such as those that produce copious trichomes, mucilage, or have a thick cuticle. Unlike floral dip, floral bud injection bypasses these defenses. For these reasons, we will primarily be using the floral bud injection method going forward.

As with any stable transformation method in plants, there are still disadvantages to the jab infiltration method. Transformation efficiency is still very low, and some genotypes/accessions, particularly inland annuals, are recalcitrant to it (for example, LMC could not be transformed by floral dip or floral bud injection, even when several parameters were modified). Especially for applications like CRISPR/Cas9 gene editing, where editing efficiency is compounded with transformation efficiency, this method will likely only work well with the most amenable genotypes.

### Using these methods to identify the mechanistic basis of ecological and evolutionary processes in Mimulus

Multiple species across *Mimulus* are transformable by tissue culture, floral dip, and now jab infiltration methods (Bell, 2021; Ding et al., 2020; Yuan et al., 2013; dx.doi.org/10.17504/protocols.io.8vghw3w; dx.doi.org/10.17504/protocols.io.3tkgnkw; Supplemental File 1). We emphasize that there is no one-size fits all solution for stable transformation in *Mimulus*, which has a wide array of phenotypes that could aid in or impede transformation. We recommend three avenues of improvement for transformation. First, we suggest that researchers identify the most fecund lines with the least inbreeding depression to develop as genetic models. Second, we encourage researchers who use different methods (tissue culture, floral dip, and jab infiltration) continue to refine and share their advances, as some methods work better for certain species or accessions. To our knowledge, no one has tried biolistics methods in *Mimulus*, and there are newer advances such as the use of nanoparticles and non-transgenic editing that could advance hypothesis testing in the system (Q. Liu et al., 2023; Lv et al., 2020). Third, we plan to explore the use of other delivery systems as a means to improve transformation efficiency (other strains of *Agrobacterium, Ensifer adhaerens*, *Ochrobactrum haywardense*, *Rhizobium etli*, etc.).

Combined with the natural history, ecological, evolutionary, population genetic, and genomic resources that the yellow monkeyflower community has built (Hellsten et al., 2013; Twyford et al., 2015; Vallejo-Marín et al., 2021; Wu et al., 2008), simple methods for genetic transformation will allow us to test hypotheses about reproductive isolation, local adaptation, and speciation in the *Mimulus guttatus* species complex in greater molecular detail than ever before.

## ACKNOWLEDGEMENTS

We would like to thank Lane Vitek and Cam Durant for assisting with experiments. We would also like to thank Yao-Wu Yuan, who pioneered the floral spray/vacuum infiltration method in monkeyflower, along with Janelle Sagawa, Riane Young, Brian Christensen, Toby Bradshaw, Barbara Frewen, and Dena Grossenbacher.

## AUTHOR CONTRIBUTIONS

L.E.S. designed and conducted all transgenic experiments. D.B.L. obtained funding and materials for the research. L.E.S. and D.B.L. wrote the manuscript.

