## Supplemental File 1 for "Just add water: A simple floral bud injection method for stable *Agrobacterium*-mediated transformation in two ecotypes of *Mimulus guttatus*"

THE “JUST ADD WATER” PROTOCOL

Materials

- *Agrobacterium tumefaciens* strain GV3101
- 1 cc syringe
- 30 G needle

Optional Materials

- Sucrose
- Silwet L-77

Steps

1. Inoculate *Agrobacterium* GV3101 containing your construct of interest into 3 mL of LB + appropriate antibiotics. Shake overnight at 200 rpm, 28 °C.

2. Decant 3 mL *Agrobacterium* culture into a flask containing 300 mL of LB + appropriate antibiotics. Shake 14-16 hours at 200 rpm, 28 °C.

3. Spin down 200-240 mL of *Agrobacterium* culture in conical tubes or bottles, 4800 g, 15 minutes, 4 °C.

4. Remove supernatant. Resuspend bacteria in 300 mL DI water^1^. Note: The bacteria will no longer have a nutrient source; **proceed immediately to the next step**.

5. Using a 1 cc syringe with 30 G needle, inject young floral buds (calyx still closed or mostly closed, no or very little visible corolla) with the *Agrobacterium* solution. Inject the base of the bud, aiming for the ovary, until the calyx darkens and the bacterial solution begins to escape the bud tip. Do not inject the same bud more than once to reduce tissue damage.

6. Return plants to their normal growing conditions.

7. Self-pollinate the flowers that were injected (there are obvious injection sites on most infiltrated buds). Note: You can keep un-infiltrated plants or use flowers from an un-infiltrated branch as pollen donors.

8. Collect fruits and plant seeds on soil. When seeds start to germinate, spray with herbicide appropriate to the construct every day or every other day^2^. Transformed seedlings will be conspicuously large and healthy in a lawn of yellowed and dying untransformed seedlings. Note: the inland annual plants grow very rapidly, so be sure to spray with herbicide immediately upon germination. Once the plants reach a certain size, they are more resistant to the herbicide.

^1^Optional in perennial *M. guttatus*: Add 24 uL Silwet L-77 (final concentration 0.01%) to the resuspension solution. Optional in annual *M. guttatus*: Add 7.5-30 g sucrose (final concentration 2.5-10% w/v) to the resuspension solution.

^2^We use 1:1000 BASTA (glufosinate-ammonium). Seeds can also be screened on agar plates with the appropriate antibiotics (e.g., hygromycin), as in *Arabidopsis*.
